# A multi-step filtering pipeline for human read removal enhances detection of *Fusobacterium* in WGS datasets with immunohistochemical confirmation in mucinous rectal cancer

**DOI:** 10.1101/2025.11.24.690231

**Authors:** Mariia Frolova, Barry Maguire, Heiko Duessmann, Menatallah Rayan, Caoimbhe Burke, Hesham M. Korashy, Arman Raman, William Gallagher, John P. Burke, Simon J. Furney, Jochen H. M. Prehn

## Abstract

The study of tumour-associated microbiomes using whole-genome sequencing (WGS) has attracted considerable attention, but microbial signal detection remains controversial due to host contamination and methodological artefacts. As the necessity of human-read removal becomes increasingly evident, many groups now include this step in their data pre-processing workflows. In this work, we introduce an open-source tool designed for rigorous host-read removal and apply it to the reanalysis of WGS data from ten mucinous rectal adenocarcinoma cases originally published by Reynolds et al. The workflow integrates k-mer-based classification (Kraken2), quality and adapter trimming (Trim Galore), vector filtering (BBDuk/UniVec_Core), and duplicate-removal (FastUniq). After reducing data complexity, a multi-aligner, multi-reference approach (BWA-MEM/GRCh38, Bowtie2/T2T-CHM13, Minimap2/HPRC v1.1) removes remaining host sequences, collectively eliminating more than 99.9% of human-derived reads. Although the additional alignment steps eliminated only a small fraction of total reads, they consistently removed millions of residual sequences per sample, underscoring the importance of rigorous filtering in datasets where non-human reads are a small minority. Taxonomic profiling with PathSeq and MetaPhlAn revealed reproducible enrichment of *Fusobacterium* species in tumour versus matched normal tissues, with stricter filtering reducing overall microbial signal compared to prior results. Cross-validation with immunofluorescence analysis using pan-*Fusobacterium* (detecting both *F. animalis* and *F. nucleatum*) and *F. nucleatum*-specific antibodies showed a strong concordance with *Fusobacterium* subspecies detected by WGS. Compared to the unfiltered analysis, host depletion markedly reduced artificial microbial signals in normal samples while preserving tumour-associated *Fusobacterium*, resulting in a more reliable microbial profile.

**Importance:** We developed an open-source tool that enables rapid removal of human-derived sequences and applied it to rectal cancer WGS data. This approach reduced false microbial signals while preserving true tumour-associated *Fusobacterium*, and we confirmed these findings in tissue using immunofluorescence staining. Our method provides a more reliable foundation for studying tumour–bacteria interactions.

## Introduction

The study of tumour microbiomes using whole genome sequencing (WGS) has attracted significant attention as bacteria have been found in various types of solid tumours including colon, pancreatic, breast cancers [1-2]. These discoveries have sparked growing interest in the potential roles of bacteria in tumour development, progression, and response to therapy. However, the validity of microbial signals in tumour tissue remains debated.

One of the most widely publicized studies in this area, published by Poore *et al*. in 2020, claimed that microbial DNA can be detected in various tumour types using data from The Cancer Genome Atlas (TCGA) [3]. However, the study was formally retracted [4] as independent investigations found serious methodological shortages. Reanalysis by Gihawi and Salzberg [5] showed that the main cause of incorrect taxonomic assignments was insufficient removal of human reads. Moreover, bacterial reference genomes can contain human sequence insertions, leading misclassification [6-7]. These findings have underscored to the research community the necessity of rigorous host read removal, and most recent studies now implement careful and comprehensive human read filtering as a standard step [8-9]. Given these challenges, it is critical to apply strict quality control and human read filtering before interpreting the presence of microbial sequences in tumour WGS data. To validate computational results, it is important to compare microbial abundance estimates obtained from WGS-based profiling with other techniques, such as immunodetection. While sequencing provides quantitative taxonomic resolution, immunostaining provides spatial and morphological context in addition to confirming these signals. Such cross-comparison is particularly important when studying closely related subspecies, where read misclassification and background noise may otherwise confound computational results. A recent study demonstrated that *F. animalis* is the predominant *Fusobacterium* subspecies detected in more colorectal tumour samples compare with *F. nucleatum* [10]. These findings highlight the importance of distinguishing between different *Fusobacterium* subspecies, as only *F. animalis* showed a prognostic impact, underscoring the need to separate their contributions to colorectal cancer development and progression. We here describe a multi-step filtering pipeline for human read removal that enhances the detection of *Fusobacterium* in WGS datasets, and confirm the presence of *Fusobacterium* subspecies in mucinous rectal cancers.

## Materials and Methods

### 1. Raw read analysis

We re-analyzed the WGS dataset published by Reynolds *et al*. [11], which includes ten cases of mucinous rectal adenocarcinoma with matched tumour and adjacent normal tissues. Reads were processed with HumanFilt (v1.0.0), a workflow developed in-house and distributed via Bioconda, configured exclusively for paired-end Illumina data (https://github.com/jprehn-lab/humanfilt). The pipeline (i) pre-filters human reads with Kraken2 against a curated database derived from GRCh38 and T2T-CHM13 reference genomes; (ii) performs adapter and quality trimming with Trim Galore, discarding reads <20 bp and bases with Phred <20; (iii) collapses duplicate read pairs with FastUniq; (iv) removes sequencing-vector contamination using BBMap bbduk.sh against the UniVec_Core database (August 2024 release; k=27, hdist=1); and (v) applies stepwise host-alignment filtering, retaining only pairs unmapped at each stage: first to GRCh38 with BWA-MEM, then to T2T-CHM13 with Bowtie2, and finally to the HPRC v1.1 pangenome with Minimap2. On first use, HumanFilt (v1.0.0) automatically downloads and caches all required references and databases from our Zenodo repository (DOI 10.5281/zenodo.17020482), including GRCh38, T2T-CHM13, HPRC v1.1, UniVec_Core, and the curated human Kraken2 database. All tools were obtained through Conda (bioconda and conda-forge). The principal tools and their versions listed in Supplementary Table S0.

### 2. Taxonomic analysis using PathSeq and MetaPhlAn

Filtered non-human reads were subjected to taxonomic profiling with two complementary methods. PathSeq (GATK v4.6.1.0) [12] was run against a curated *Fusobacterium* reference database that had been pre-screened to exclude potential human insertions using a sliding-window strategy (150 bp windows, 75 bp steps) derived from GRCh38, T2T-CHM13 v2.0, and HPRC v1.1, aligned against Fusobacterium genomes with Bowtie2. In parallel, MetaPhlAn (mpa_vJan25_CHOCOPhlAnSGB_202503) [13] was applied with default parameters to provide independent estimates of microbial abundance. Relative abundances of *Fusobacteriota* phylum members and selected clades (*F. nucleatum* and *F. animalis*) were quantified in tumour and matched normal samples. Results from our re-analysis were compared with the original findings of Reynolds *et al*. to evaluate the impact of stricter human-read filtering on microbial signal detection.

### 3. Histological slice preparation, staining, and image analysis

Formalin-fixed paraffin-embedded (FFPE) tumour sections from primary tumour sections of mucinous rectal cancer patients following resection at the Beaumont RCSI Cancer Centre (BRCC) were obtained. Institutional ethical approval was granted by the Beaumont Research and Ethics Committee (Reference 21/98). Sections were cut (5 µm), baked, and mounted on individual slides. Prior to staining, the slides were baked at 60 °C overnight. The Leica Bond RXm platform (Leica Biosystems, Nussloch, Germany) was used henceforth for slide preparation and staining. Sections underwent deparaffinisation, rehydration, permeabilization in 0.3% Triton X-100 in PBS for 10 minutes, and heat-mediated epitope retrieval using Bond ER1 (Citrate-based, pH 6.0) and ER2 (Tris-based, pH 9.0) buffers. Slides were then incubated in blocking solution (10% donkey serum, 3% BSA in PBS) for 1 hour at room temperature. After blocking, the slides were immersed in PBS and exposed to a multi-spectrum LED for 48 hours to photoirradiate the tissue. Nuclei were stained with DAPI 1 µg/mL for 15 minutes and the Cell DIVE platform (Leica Microsystems, Wetzlar, Germany) was used to perform 2X whole slide imaging and tissue selection, 10X whole tissue imaging and region selection, and 20X autofluorescence imaging. Following that, during the first round of staining, slides were incubated for 1 hour with a commercial antibody (ANT0084 1:100; Diatheva, Cartoceto, Italy) raised against *F. nucleatum* subsp. *nucleatum* 25586 which preferentially detects *F. nucleatum*. Following primary antibody staining, slides were washed 3 times for 30 seconds in Bond Wash buffer then incubated for 1 hour at room temperature with the Alexa Fluor 647 donkey anti-rabbit antibody (Abcam, Cambridge, UK). Subsequently, the slides were stained using a pan-*Fusobacterium* antisera (1:100) raised against purified total membrane proteins from 11 stains of *Fusobacteria* (*F. nucleatum* subsp. *nucleatum* 23726, *F. nucleatum* subsp. *animalis* 7_1, *F. nucleatum* subsp. *polymorphum* 10953, *F. nucleatum* subsp. *vincentii* 49256, *F. periodonticum* 2_1_31, *F. varium* 27725, *F. ulcerans* 49185, *F. mortiferum* 9817, *F. gonidiaformans* 25563, *F. necrophorum* subsp. *necrophorum* 25286, *and F. necrophorum* subsp. *funduliforme* 1_1_36S) [14] recognizing both *F. nucleatum* and *F. animalis* [15], [16], followed by an Alexa Fluor 555 donkey anti-rabbit antibody (Abcam, Cambridge, UK). Nuclei were counterstained with DAPI and 20X biomarker imaging was undertaken between each staining round. Digital images were processed and analysed using QuPath (v0.6.0) for tissue annotation, background correction, and visualisation. To confirm the appearance of *Fusobacterium* morphology, slides were then coverslipped and imaged using the LSM 980 Airyscan 2 confocal microscope (Carl Zeiss, UK) using a 63x 1.4NA oil immersion objective. Confocal z-stacks of optical slices of *Fusobacterium*-positive regions were at 150 nm intervals with a pixel size of 35 x 35 nm using the 4Y-mode. Images were processed with ZEN software (Carl Zeiss, UK).

### 4. Cell and bacterial co-culture, and staining and imaging

*Fusobacteria* were cultured overnight in fastidious anaerobe broth (FAB; Neogen) at 37ºC under 0.1% O_2_ using an anaerobic chamber (Whitley A35; Don Whitley Scientific). To remove dissolved oxygen, FAB and PBS were autoclaved and allowed to equilibrate overnight in the anaerobic chamber prior to use. Cancer cells (MDA-MB-468) were seeded in cover glass bottom cell culture dishes (μ–dish #81156, IBIDI, Germany). Once desired confluency was reached (70-80%), cells were infected with an optimised multiplicity of infection (MOI) of 1:125. After 12 hours of co-culture, extracellular bacteria were removed by washing twice with PBS and subsequent incubation for one hour in DMEM/F-12 containing 200 μg/ml metronidazole and 300 μg/ml gentamicin. Cells were then washed and fixed with 4% formalin for 20 min at RT and stored at 4ºC.

Immunocytochemical (ICC) staining of cells was undertaken. First, cells were permeabilised using a buffer containing 1% BSA, 2% horse serum and 0.1% Triton in PBS for 30 minutes at room temperature. Cells were washed three times with PBS for 3 minutes each. Antibody solutions were prepared in the ICC buffer with either the commercial *F. nucleatum* antibody (1:100) (ANT0084) or the pan-*Fusobacterium* antisera (1:100) along with a Pancytokeratin antibody (1:100) (Abcam, Cambridge, UK) and incubated for 1 hour at room temperature. After an further three 3-minute washes with PBS, cells were stained with a cocktail solution of Alexa Fluor 555 goat anti-rabbit antibody (1:1000) (Abcam, Cambridge, UK) and Alexa Fluor 647 donkey anti-mouse antibody (1:1000) (Abcam, Cambridge, UK) for 1 hour at room temperature. Following a subsequent wash, cells were incubated with Hoechst 1mg/ml for 30 minutes at room temperature. After a final round of PBS washes, cells were imaged on a confocal super-resolution microscope (LSM 980 Airyscan 2, Carl Zeiss UK) using a 63x 1.4NA oil immersion objective. Stacks of optical slices were taken at an interval of 150 nm with a pixel size of (35 nm)^2^ using the 4Y-mode. Imaging data was prepared for visualisation using the same range of the histogram for each staining in ZEN (ZEN, Carl Zeiss UK).

## Results

### 1. Filtering of human reads

We implemented a stepwise pipeline to remove host and contaminant sequences from mucinous rectal cancer WGS data (Figure 1). The workflow HumanFilt-1.0.0 removed human and vector sequences, with read counts recorded at each processing step and compiled into a final summary report for each sample (https://github.com/jprehn-lab/humanfilt). Raw reads were first processed with Kraken2 [17]against a curated human database (GRCh38 + T2T-CHM13), removing 99.6% ± 0.08% of total reads per sample. Because Kraken2 is k-mer–based, adapter sequence has negligible impact at this stage, so this pass both accelerates processing and significantly reduces the dataset for subsequent steps. We then applied Trim Galore [18] (Phred ≥20; minimum length 20 bp) for adapter and quality trimming to improve mapping ability, followed by FastUniq [19] to collapse duplicate read pairs. Sequencing-vector contamination was filtered with BBDuk (BBTools v37.62) [20] against UniVec_Core (August 2024; k=27, hdist=1), typically removing ∼54,000–223,000 reads per sample. Finally, the remaining unclassified reads were aligned sequentially to GRCh38 [21] (BWA-MEM [22]), T2T-CHM13 [23] (Bowtie2 [24]), and the HPRC v1.1 pangenome [25] (Minimap2 [26]), retaining only pairs unmapped at each stage. These alignment filters removed an additional 0.23–0.38% of residual human reads ∼1.7–4.0 million per sample yielding the final high-confidence non-host set. Read counts at each stage of HumanFilt were computed with SeqKit [27]. The proportion of reads retained after each filtering step is summarized in Table 1 (normalized) and Supplementary Table S1 (not-normalized, tumour samples), Supplementary Table S2 (not-normalized, normal samples).

**Table 1.** Read processing summary for mucinous rectal cancer WGS data.

| Case ID | Total Reads (M) | Kraken2 Unclassified (T) |  | Trimmed Reads (T) |  | Vector & Duplicate Reads Removed (T) |  | Unaligned Reads (GRCh38 +T2T+HPRC) (T) |  |
| --- | --- | --- | --- | --- | --- | --- | --- | --- | --- |
| A | 1010 | 4484 | 0.44% | 4422 | 0.44% | 4328 | 0.43% | 815 | 0.08% |
| B | 610 | 1900 | 0.31% | 1853 | 0.3% | 1799 | 0.29% | 146 | 0.02% |
| C | 1013 | 2832 | 0.28% | 2759 | 0.27% | 2680 | 0.26% | 295 | 0.03% |
| D | 1011 | 4583 | 0.45% | 4516 | 0.45% | 4423 | 0.44% | 944 | 0.09% |
| E | 1007 | 3083 | 0.31% | 3021 | 0.3% | 2921 | 0.29% | 252 | 0.02% |
| F | 938 | 2738 | 0.29% | 2655 | 0.28% | 2567 | 0.27% | 209 | 0.02% |
| G | 1015 | 3577 | 0.35% | 3491 | 0.34% | 3374 | 0.33% | 542 | 0.05% |
| H | 1022 | 3971 | 0.39% | 3783 | 0.37% | 3597 | 0.35% | 853 | 0.08% |
| I | 1092 | 4945 | 0.45% | 4757 | 0.44% | 4534 | 0.41% | 967 | 0.09% |
| J | 551 | 2684 | 0.49% | 2613 | 0.47% | 2512 | 0.46% | 455 | 0.08% |

**Figure 1.**
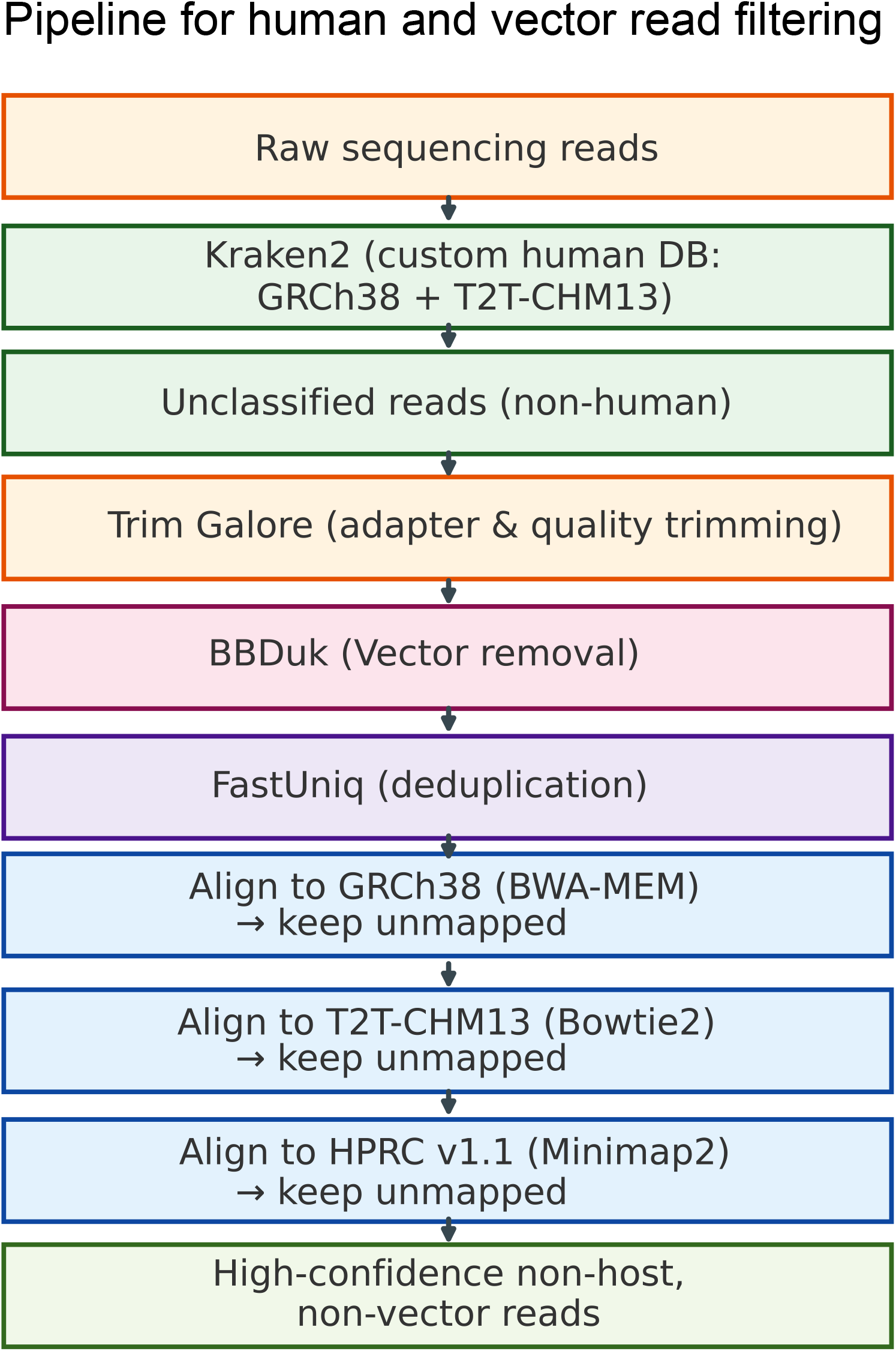
Workflow for filtering human and vector reads from WGS data.

### 2. Comparison of Fusobacteriota and F. nucleatum abundance between studies

Before constructing the PathSeq [12]*Fusobacterium* database, we assessed all reference genomes included in the genus for potential human sequence insertions. Using the sliding-window strategy described by Poore *et al*. [28], 150 bp sequences were generated at 75 bp intervals from GRCh38, T2T-CHM13v2.0, and the HPRC contig assembly v1.1, excluding fragments shorter than 75 bp or containing more than100 ambiguous bases. These sequences were aligned to the *Fusobacterium* reference genomes using Bowtie2. No human-derived sequences aligned to any *Fusobacterium* genome, confirming that the genus-wide references are free from human DNA contamination.

After implementing multiple filtering steps to remove human reads within our analysis pipeline, we compared *Fusobacteriota* phylum read abundance between Reynolds *et al*. and our current reanalysis across ten matched tumour and normal tissue samples (Cases A–J) (Figure 2). In both studies, *Fusobacteriota* were detected by PathSeq, and tumour samples consistently exhibited higher levels than their matched normal tissues (Figure 2A). In the results from Reynolds *et al*., the abundances were notably higher, particularly in Cases G–J (up to 23.56%). By contrast, our re-analysis showed lower overall percentages across both tumour and normal samples, suggesting stricter filtering. Tumour enrichment remained evident in several cases (e.g., Cases C, E, G, H, I, J), though the magnitude of this difference was reduced, and normal samples displayed substantially lower background levels compared to the finding of Reynolds *et al*..

**Figure 2.**
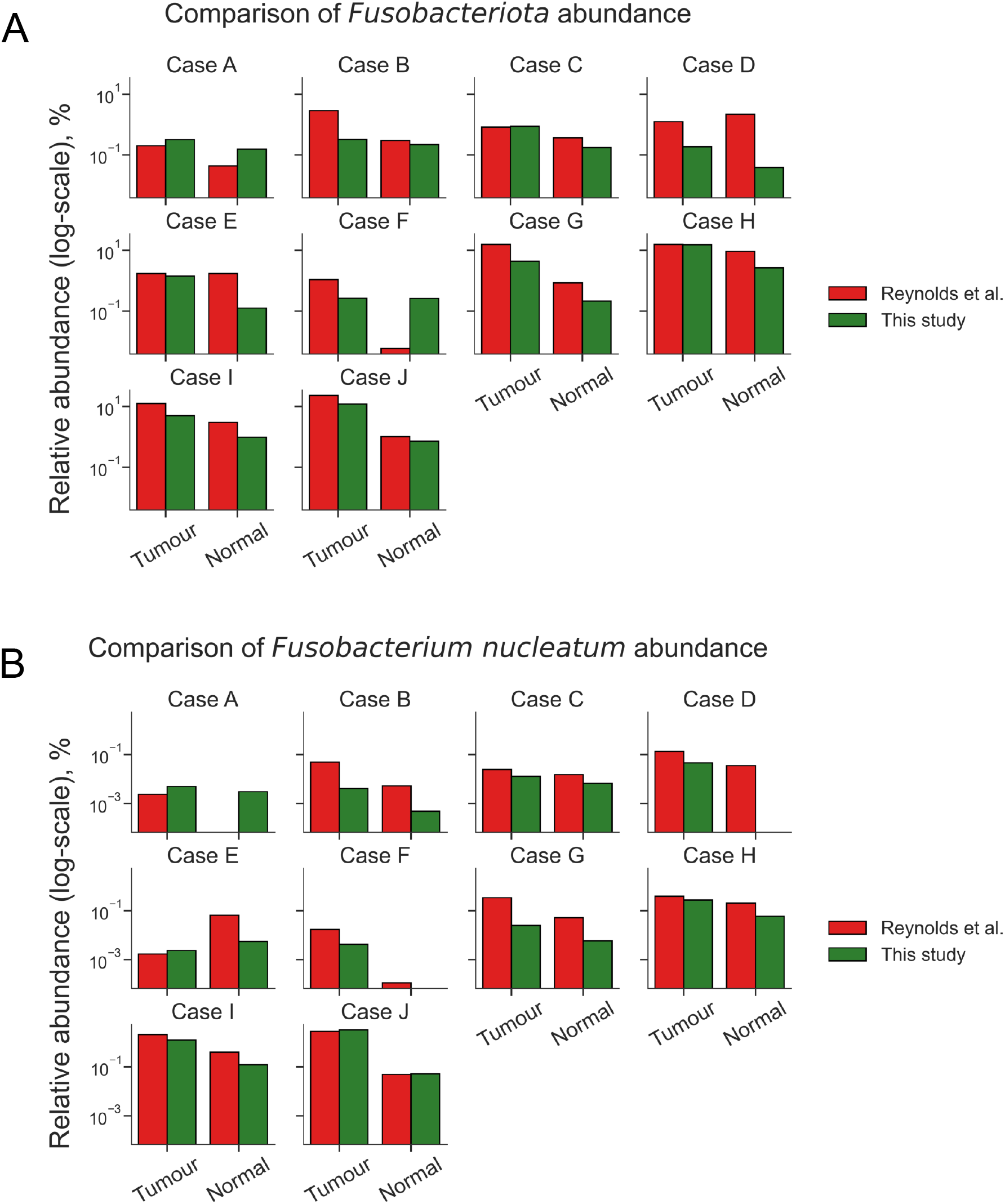
Relative abundance (%) of (A) *Fusobacteriota* phylum and (B) *F. nucleatum* species in matched normal and tumour tissue samples from mucinous adenocarcinoma of the rectum. Barplots compare read percentages from Reynolds et al.’s analysis and the current study, in which human reads were removed using our bioinformatics pipeline.

We further examined the relative abundance of *F. nucleatum*. In both analyses, *F. nucleatum* was detected by PathSeq and enriched in tumours compared with normal tissues (Figure 2B). In the analysis of Reynolds *et al*., the highest levels were observed in Cases I and J (2.08% and 2.83%, respectively). In our re-analysis, overall abundances were lower, but enrichment remained evident, particularly in Case J (3.29%) and Case I (1.25%). Notably, for Case I our relative abundance was higher than previously reported, because the denominator of total reads after filtering differed, leading to a higher proportional value. Normal tissues in our dataset showed markedly reduced background levels—often near zero or undetectable— especially in Cases D and F. Together, these findings demonstrate that *Fusobacteria*, including *F. nucleatum*, are reproducibly detected in colorectal tumour samples by PathSeq, while our stricter filtering pipeline reduces background noise and likely yields more accurate abundance estimates.

Detailed relative abundance values for *Fusobacteriota* and *F. nucleatum* for all samples are provided in Supplementary Tables S3 and S4, respectively.

### 2. Correlation between WGS-based clade detection and α-Fusobacteria immunostaining

To determine whether WGS-based detection of Fusobacterium reflects bacterial presence within tumour tissue, we compared the PathSeq [12]and MetaPhlAn [13] outputs with immunostaining using the pan-*Fusobacterium* and *F. nucleatum*-specific (ANT0084) antibodies (Figure 3). PathSeq provides sensitive species-level detection and absolute read counts (RPM), while MetaPhlAn reports marker-based relative abundances with higher specificity, which increases confidence in species assignment and in determining the dominant *Fusobacterium* clade. Using both reduces false positives and false negatives and enables direct comparison with staining results.

**Figure 3.**
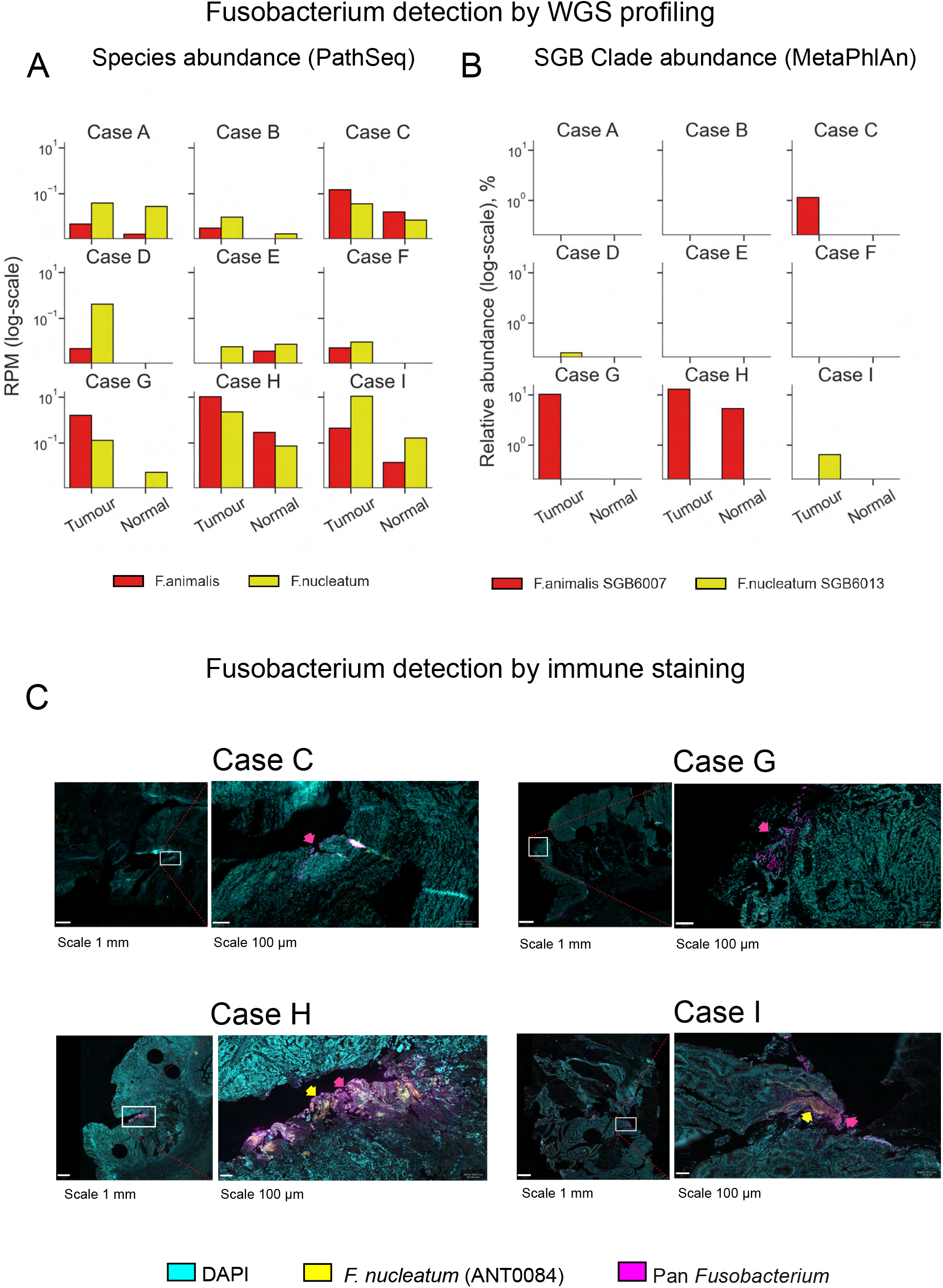
Detection of *F. nucleatum* and *F. animalis* by sequencing and immunostaining. (A) Normalized read counts (RPM, log-scale)) of *F. nucleatum* and *F. animalis* across tumour and normal samples as determined by PathSeq. (B) Relative abundance (%, log-scale) of the same species as determined by MetaPhlAn, where *F. animalis* was identified as SGB6007 and *F. nucleatum* as SGB6013 clades. (C) Immunostaining results using two rabbit-derived *Fusobacteria* antibodies: the pan-*Fusobacterium* antibody (detecting both *F. nucleatum* and *F. animalis*) and the commercial *F. nucleatum*-specific ANT0084 antibody.

In the stained samples, MetaPhlAn detected only *F. animalis* (SGB6007) and *F. nucleatum* (SGB6013), while other *Fusobacterium* clades (SGB6011, SGB6001, SGB6014, SGB5985, SGB5986) were not identified. This is relevant because the pan-*Fusobacterium* antibody can detect multiple *Fusobacterium* species. Although low-level reads from other species were detectable by PathSeq (Supplementary Table S7), they were not dominant and are unlikely to explain the observed staining patterns with the pan-*Fusobacterium* antibody. Thus, staining only with the pan-*Fusobacterium* antibody suggests the presence of *F. animalis*, staining only with ANT0084 indicates *F. nucleatum*, and positive staining with both antibodies supports the presence of both species.

Overall, we observed a strong correspondence between WGS results and antibody staining patterns (Table 2, Figure 3).

**Table 2.** Comparison of WGS-based *Fusobacterium* detection and immunostaining results.

| Case ID | Dominant species by WGS | PathSeq (RPM) | MetaPhlAn (%) | Pan-Fusobacterium staining | F. nucleatum-specific (ANT0084) staining | Interpretation |
| --- | --- | --- | --- | --- | --- | --- |
| Case C | *F. animalis* | 0.15 | 1.13 | + | – | *F. animalis* |
| Case G | *F. animalis* | Fa 1.6 / Fn 0.13 | Fa 10.2 / Fn 0 | + | + | *F. animalis*-predominant |
| Case I | *F. nucleatum* | Fa 0.44 / Fn 10.9 | Fa 0 / Fn 0.65 | + | + | *F. nucleatum*-predominant |
| Case H | *F. animalis* | 10.5 | 12.7 | + | – | *F. animalis* |

Case C and Case H both provide strong examples of *F. animalis* positive, *F. nucleatum* low tissue as validated by both WGS and immunostaining. In these cases, species quantification through WGS showed clear dominance of *F. animalis*, while *F. nucleatum* was detected only at background or very low levels (Table 2, Supplemental Tables S5, S6). Immunostaining results corresponded, showing positive staining with pan-*Fusobacterium* antibody but little to no staining with the *F. nucleatum*-specific antibody (Figure 3C). This pattern suggests that while small counts of *F. nucleatum* may be present, they are below the threshold for antibody detectability and unlikely to represent a major component of the microbial community in these samples.

For the F. *nucleatum* positive category, Case I serves as an example, showing high levels of *F. nucleatum* by WGS (Table 2, Supplementary Tables S5, S6) and strong staining with both the *F. nucleatum*-specific and pan-*Fusobacterium* antibodies (Figure 3), indicating *F. nucleatum* predominance within the tissue sections. Although the pan-*Fusobacterium* antibody can also detect *F. animalis*, the absence (or only trace levels) of *F. animalis* reads in the WGS data supports that the observed staining primarily reflects *F. nucleatum*.

In contrast, Case G showed high *F. animalis* abundance and a low but detectable *F. nucleatum* signal by WGS (Table 2, Supplementary Table S5). Both the pan-*Fusobacterium* and *F. nucleatum*-specific antibodies exhibited positive immunostaining (Figure 3); however, the WGS data indicated a predominance of *F. animalis*, suggesting that the tissue was primarily colonized by this species.

Taken together, we observed a strong concordance of sequencing and antibody-based detection of *Fusobacteria*.

### 4. Validation of species-specific immunostaining using controlled infections

To validate species-specific immunostaining, commercial strains of *F. nucleatum* (ATCC 25586) and *F. animalis* (ATCC 51191) were separately co-cultured with MDA-MB-468 breast cancer cells. Infected and control cells were stained with the pan-*Fusobacterium* antiserum and a commercial *F. nucleatum-*specific antibody (ANT0084), alongside pancytokeratin and Hoechst nuclear stain. High-resolution confocal microscopy (Zeiss LSM 980 Airyscan 2) was employed to acquire optical stacks.

Cells infected with *F. animalis* displayed clear intracellular fluorescence with the pan-*Fusobacterium* antiserum (Figure 4A, bottom middle), but no staining with ANT0084 (Figure 4A, top middle), confirming that the pan-Fusobacterium reagent detects *F. animalis* while ANT0084 is specific to *F. nucleatum*. Conversely, *F. nucleatum*–infected cells exhibited robust staining with both the pan-*Fusobacterium* antiserum (Figure 4A, bottom left) and ANT0084 (Figure 4A, top left). Uninfected control cells stained with ANT0084 were negative (Figure 4A, top right), while those stained with the pan-*Fusobacterium* antiserum showed only very weak background signals (Figure 4A, bottom right), indicating high specificity.

**Figure 4.**
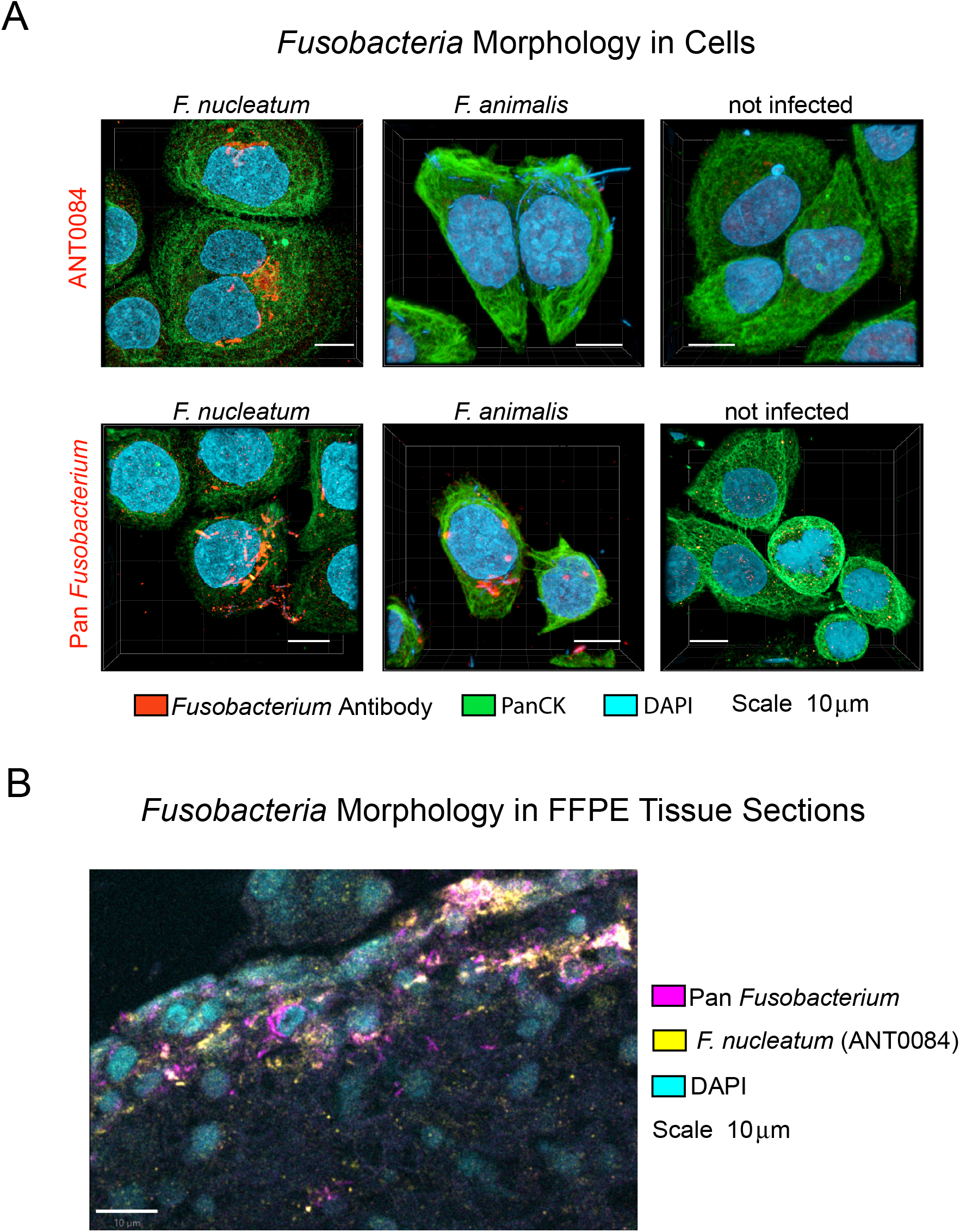
Immunofluorescence validation of *Fusobacterium* in MDA-MB-468 cells. Cells where infected with *F. nucleatum* (left column) *F. animalis* (middle column), and not infected (right column). They were stained with the commercially available ANT0084 (top row) and pan-*Fusobacterium* antiserum (D. Slade, bottom row). Shown are maximum intensity projections of z-Stacks, scalebar is 10 μm in x and y. Imaging parameters. Zeiss LSM 980 Airyscan 2, 63×/1.4 NA; voxel size 35 × 35 × 150 nm^3^ (dx, dy, dz); estimated resolution Δx = Δy ≈ 120 nm, Δz ≈ 400 nm; field of view (x) ≈ 44.47 µm. Images acquired as optical section stacks.

Taken together, these data confirm that the pan-*Fusobacterium* antiserum recognizes multiple *Fusobacterium* species including *F. nucleatum* and F. *animalis*, whereas ANT0084 is a reliable species-specific antibody for *F. nucleatum*. This dual approach supports the WGS-based assignments of *Fusobacterium* species in tissue analyses.

To extend these findings to clinical material, we included a representative confocal image illustrating *Fusobacterium* morphology in FFPE tissue sections (Figure 4B). The same pan- *Fusobacterium* antiserum and *F. nucleatum*-specific antibody (ANT0084) were applied to the sections, together with DAPI nuclear staining. A similar pattern of staining was observed in the tissue as in cell culture, with *F. nucleatum* positive regions displaying overlapping signals from both antibodies. The pan-*Fusobacterium* antiserum produced a clear positive signal in infected regions but also exhibited weak background staining in noninfected tissue, consistent with the observations in cell culture.

## Discussion

With growing interest in tumor-associated microbiomes, WGS interpretation remains vulnerable because non-human reads constitute a small minority and taxonomic profiling tools can misclassify human reads as bacterial. To address this, we developed a workflow that prioritizes high specificity and computational efficiency, carefully excluding human reads to minimize errors in subsequent steps. We implemented a pre-screen-then-align strategy: first applying rapid k-mer–based host depletion, then performing quality and contaminant clean-up and ultimately executing sequential alignments to complementary human references to eliminate residual host signal.

Kraken2 pre-screening against a curated human k-mer database efficiently eliminates the overwhelming bulk of human sequence and reduces data volume for all downstream operations. Because Kraken2 relies on exact k-mer matches, adapter sequence contributes no informative k-mers and typically has negligible impact at this stage. Any minor loss of k-mer informativeness caused by adapter-rich reads is corrected by the subsequent trimming and alignment stages. We therefore trim and quality-filter after pre-screening, collapse duplicate pairs, and remove vector sequences to improve mapping capability precisely where aligners are most sensitive—adapters, low-quality tails, and technical contaminants. Residual host signal is then removed with a multi-aligner, multi-reference strategy that retains only read pairs unmapped at each stage.

Different aligners work differently: they use distinct strategies to find matches, score them, and backtrack. As a result, each aligner catches some human reads that another misses, even on the same reference genome. To avoid overloading the workflow we maintain aligner diversity but assign each tool to a separate human reference optimized for its mapping profile.

Our design leveraged three aligners—BWA-MEM, Bowtie2, and Minimap2 - each suited to different genomic contexts. BWA-MEM, optimized for linear reference mapping, performs robustly when aligned to GRCh38, the traditional human genome assembly [29]. In contrast, Bowtie2, with its backtracking algorithm and ability to handle mismatches and gapped alignments, is particularly effective for mapping reads within highly repetitive regions, such as centromeres and telomeres, which are fully represented in the T2T-CHM13 assembly [30], [31]. We selected Minimap2 for alignment against the HPRC v1.1 pangenome reference because its algorithm is particularly well suited to the structural and haplotype complexity represented in this assembly [32], [33]. By combining these tools with complementary references, we maximized the coverage and efficiency of human-read removal across diverse genomic contexts. While Kraken2 removed the vast majority of human reads, our results show that the subsequent alignment-based filtering steps were also essential. Although these accounted for only a small percentage of the total reads, they consistently removed millions of sequences per sample, underscoring the importance of stringent filtering in low-biomass datasets where even modest levels of host contamination can distort microbial profiles.

Compared with previously published unfiltered data [11], our conservative workflow demonstrates clear benefits: although absolute microbial signals are reduced, especially in matched normal samples, the substantial decrease in background noise enhances confidence in detecting true tumor-associated microbiota. Our findings also reveal limitations of relying solely on relative abundance metrics in low-biomass samples, where differences in denominators after human-read filtering can artificially inflate apparent taxon prevalence. For instance, in Case J, *F. nucleatum* (Figure 2) showed higher relative abundance than previously reported despite similar or lower absolute read counts. This compositional bias, typical of metagenomic tools like MetaPhlAn, can be mitigated by incorporating absolute quantification measures such as reads per million (RPM) calculated before host-read filtering, providing a more accurate estimate of bacterial signal.

Our immunohistochemical staining showed strong agreement with WGS data, providing robust spatial and morphological confirmation of bacterial species identified by sequencing. Recent studies suggest that *F. animalis* may play a more aggressive role in colorectal cancer progression compared to *F. nucleatum* [34], [35], so calling subspecies correctly matters. Our findings emphasize the importance of careful removal of human contamination and accurate quantification of bacterial reads in WGS data are essential for reliable diagnosis and differentiation of *Fusobacterium* subspecies in colorectal cancer.

## Acknowledgements

We are deeply grateful to the patients who provided samples, making this research possible.

This work was funded by the Higher Education Authority North South Collaboration grant (AICRI-START) to JHMP and WG, an RCSI MD StAR Fellowship to BM, and an RCSI International PhD Fellowship to MR.

We also thank the Information Technology Department at the Royal College of Surgeons in Ireland and the DJEI/DES/SFI/HEA Irish Centre for High-End Computing (ICHEC) for the provision of computational facilities and support.

